# Does method of field preparation affect survival and growth of juvenile Chinook salmon in agricultural floodplains?

**DOI:** 10.1101/2024.02.02.578246

**Authors:** Rachelle L. Tallman, Alexandra N. Wampler, Gabriel P. Singer, Carson A. Jeffres, Dennis E. Cocherell, Jordan Colby, Nann A. Fangue, Robert A. Lusardi, Andrew L. Rypel

## Abstract

There is growing interest to integrate conservation initiatives into agricultural practices using a reconciliation ecology framework. In California’s Central Valley, one approach to improving crucial nursery habitat for threatened and endangered fish species is by re-creating floodplain habitats through the inundation of agricultural fields during the non-growing season. We conducted a series of field experiments in agricultural floodplains (winter-flooded rice fields that were once natural floodplains) to explore whether different field preparation methods enhanced growth and survival of floodplain-dependent fish species. Approximately 8,000 juvenile fall-run Chinook Salmon (*Oncorhynchus tshawytscha*) were reared for at least 28 days on eight, 0.2 ha experimental fields. Each experimental field represented one of four treatments (two replicates per treatment): addition of large wood, addition of in-field canals, addition of both large wood and in-field canals, and control-where no additional field preparations were implemented beyond standard post-growing practices: i.e., chopping rice into stubble, baling and removing excess rice straw, and discing the field with a single pass by plowing rice straw into the ground to promote decomposition. We found no significant difference in water temperature, fish growth or fish survival among habitat treatments. Across treatments, survival ranged from 50.1%-78.3% and averaged 65.75% (⁺/- 7.89% SE). These findings suggest agricultural floodplains require no additional modifications to promote fish growth and survival. Results illustrate the benefits of integrating working landscapes with conservation initiatives by creating more accessible and beneficial habitat for native fishes.

## 1 INTRODUCTION

Only 0.5% of the world’s water supply is readily available freshwater found on the surface of lakes and rivers (Gleick, 1998; la Rivière, 1989), making freshwater ecosystems some of the most vulnerable habitats across the globe (Cooke et al., 2016; Dudgeon et al., 2006; Moyle & Leidy, 1992; Richter et al., 1997). Large-scale land alterations often result in habitat loss which disrupts freshwater biodiversity at all scales (Carpenter et al., 2011). Habitat loss and degradation are considered the greatest threats to freshwater vertebrates (Collen et al., 2014). This is because freshwater ecosystems are insular environments comprised of species that are highly endemic and often have narrow geographic ranges (Dudgeon et al., 2006; Strayer & Dudgeon, 2010). To maintain freshwater species, restoration is viewed as a key method to improving aquatic habitats for sensitive species (National Research Council, 1992). Restoration is used to promote native species by converting habitat back to some historical state (often pre-European colonization), however this is not always possible given the limited amount of land available and the extent of degradation (Rosenzweig, 2003a, 2003b). Instead, reconciliation ecology concepts may be used to promote conservation of freshwater species by re-establishing ecological functions within anthropogenically modified landscapes (Dudgeon et al., 2006; Moyle, 2014; Rosenzweig, 2003b; Shabtay et al., 2018)

In the last two decades, salmon populations along the western coast of the United States have reached historically low abundances (Kendall et al., 2017; Moyle et al., 2017; Zimmerman et al., 2015). Salmon serve as important linkages between the terrestrial and marine environments (MacAvoy et al., 2009). Their bodies contain marine-derived nutrients (MDN) obtained during their ocean life-history stage. MDN are subsequently released into nutrient poor streams and rivers through carcass biomass and excretion (MacAvoy et al., 2009). Pacific salmon were once abundant throughout western North America, with many populations exhibiting diverse life-history traits (Moyle, 2002). These traits provided important stock resilience against stochastic perturbations (Carlson & Satterthwaite, 2011; Katz et al., 2017; Moyle, 2002; Yoshiyama et al., 1998). Over time, riverine habitat loss associated with dams, levees, climate change, flow alteration, and other anthropogenic impacts have reduced life-history diversity across the landscape making these populations less resilient to change (Carlson & Satterthwaite, 2011; Moyle et al., 2017).

California’s Central Valley once supported the largest commercial salmon fishery in the state and contained over 1.6 million hectares of wetland habitat (Holmes et al., 2021; Yoshiyama et al., 1998). During high precipitation events, the valley floor would inundate and create large expanses of natural floodplain habitat. The relatively warm, slower floodplain waters offered juvenile salmon refugia from piscivorous fish and extensive foraging opportunities compared to the swift moving riverine environment (Grosholz & Gallo, 2006; Jeffres et al., 2008, 2020; Sommer et al., 2001; Sommer et al., 2001). As floodwaters receded, juvenile fish returned to the river, likely with enhanced lipid reserves sufficient to complete their out-migration to the Pacific Ocean. Today only 5% of the historical floodplain habitat persists in this region (Hanak et al., 2011), representing a major loss of critically important rearing habitat previously used by juvenile salmon. Wetlands were drained and converted to agriculture and cities. Dams and levees have altered natural flow regimes which has inhibited floodplain formation and prevented fish and other aquatic taxa from accessing floodplain rearing habitat (Yarnell et al., 2015). Habitat loss and degradation has greatly contributed to the decline of Central Valley Chinook Salmon (*Oncorhynchus tshawytscha*) (Yoshiyama et al., 1998). Today the stock is largely composed of hatchery supported populations (Moyle et al., 2017; Sturrock et al., 2019; Willmes et al., 2018), while remaining floodplain habitat mainly exists within managed agricultural parcels in flood bypasses (Katz et al. 2017).

There are growing efforts within the agricultural and conservation communities to implement a reconciliation framework by modifying flooding practices on working landscapes, such as agricultural fields, for native species. In the 1990’s, this framework was successfully applied within California’s Central Valley when farmers switched from burning to flooding their fields as a straw disposal method creating beneficial habitats for local bird populations (Bird et al., 2000; Strum et al., 2013). Winter-flooded fields provided additional foraging habitat for waterfowl and supported migratory bird populations that were using the Pacific Flyway (Bird et al., 2000; Strum et al., 2013, 2013). Today, this framework is considered a successful management conservation practice because it achieved a multi-benefit land use (Bird et al., 2000). Given the success of these programs for migratory birds, conservation scientists are now interested in developing a parallel framework to promote native salmon populations.

Recent efforts to apply a reconciled approach towards salmon recovery have focused on rearing juvenile salmon on winter-flooded rice fields (hereafter called agricultural floodplains) during the non-growing season (Cordoleani et al., 2022; Holmes et al., 2021; Katz et al., 2017; Sommer et al., 2001). Previous studies found that agricultural floodplains in the Central Valley exhibit protracted water residence times, increased water temperatures, and high food productivity when compared to adjacent river channel habitats (Cordoleani et al., 2022; Corline et al., 2017; Jeffres et al., 2020; Katz et al., 2017). Jeffres et al. (2020) found that fish reared on agricultural floodplains grew 7X faster compared to individuals reared in adjacent mainstem river channels. These results suggest that agricultural floodplains can support juvenile salmon with appropriate field management.

Despite these growth benefits, uncertainty and concern remains over juvenile salmon survival once these practices are adopted at larger scales and applied under drought conditions. In this scenario, there is concern over the lack of refugia within agricultural floodplains. During a drought, these habitats may be perceived as potential traps when agricultural floodplains are completely disconnected from the mainstem of the river channel and have unfavorably warm water temperatures and low dissolved oxygen concentrations. These conditions may restrict volitional passage, leaving fish vulnerable to predation as they reside in the only wetland habitat available. In this case, a management action originally intended to promote survival, may have environmental conditions that produce the opposite effect.

Physical manipulations of agricultural floodplain habitat (e.g., large wood and canals) represent potential ways of managing agricultural floodplains for the benefit of salmon. Such efforts may be especially critical in creating a “safe-operating space” (Carpenter et al., 2017) for juvenile fish during a drought (Gaeta et al., 2014; Holmes et al., 2021). Further, field preparation and design represent one component that is easily adaptable to support juvenile salmon. A reconciled approach that is too demanding with little benefit for native fish may not be taken up voluntarily in working landscapes. Alternatively, a conservation practice that requires little-to-no extra labor may not be effective in generating the desired ecological benefit. As such, we created a study that implemented additional habitat structures using methods most likely to be applied as a conservation practice and accepted by farmers in working landscapes.

Here, we evaluate the effects of additional habitat complexity (large wood, canals, large wood and canals, control-no treatment) on juvenile Chinook Salmon growth and survival in agricultural floodplains during a drought. Previous studies indicate deeper water may reduce avian predation (Brown, 2002; Cederholm et al., 1988) and prevent thermal stress in fish by providing refuge from adverse water temperatures (Ebersole et al., 2001, 2006; Tabor et al., 2011; Torgersen et al., 2012), while large wood creates important cover against predation that might improve survival (Cederholm et al., 1997; Dolloff, 1986; Radinger et al., 2023; Sass et al., 2017, 2019). We investigated if increasing water depths via canals and the addition of large wood would provide additional refuge that supports higher survival and growth rates for fish in managed agricultural floodplains. We hypothesized that differences in survival and growth rates across treatments would be the result of differences in available cover and thermal refuge.

## 2 METHODS

### 2.1 Study Site

We conducted our study at River Garden Farms (38.84 °N, −121.73 °W), a 6070 ha agricultural parcel located 42 km northwest of the city of Sacramento, CA (Figure 1). Historically, this area was prone to flooding, however construction of nearby levees and weirs redirects water away from the Sacramento River into two nearby flood bypasses (Sutter and Yolo Bypass) when the river exceeds the height of the weir. Previous research of rearing salmon on agricultural floodplains occurred within the Sutter and Yolo Bypasses(Corline et al., 2017, 2021; Holmes et al., 2021; Jeffres et al., 2020; Katz et al., 2017; Sommer et al., 2001; Sommer et al., 2005). Instead, we chose to conduct our study outside of the bypasses to prevent any potential flooding from disrupting the experiment. Rice is the main agricultural crop harvested at this farm with smaller plots dedicated to alfalfa, sunflowers, tomatoes, and walnuts, with annual crops periodically rotated. The 2020 hydrologic year (Oct 1^st^, 2019- September 30^th^, 2020) was relatively dry, creating conditions where surrounding natural wetland habitat was limited (California Hydroclimate Reports, 2020).

**FIGURE 1.**
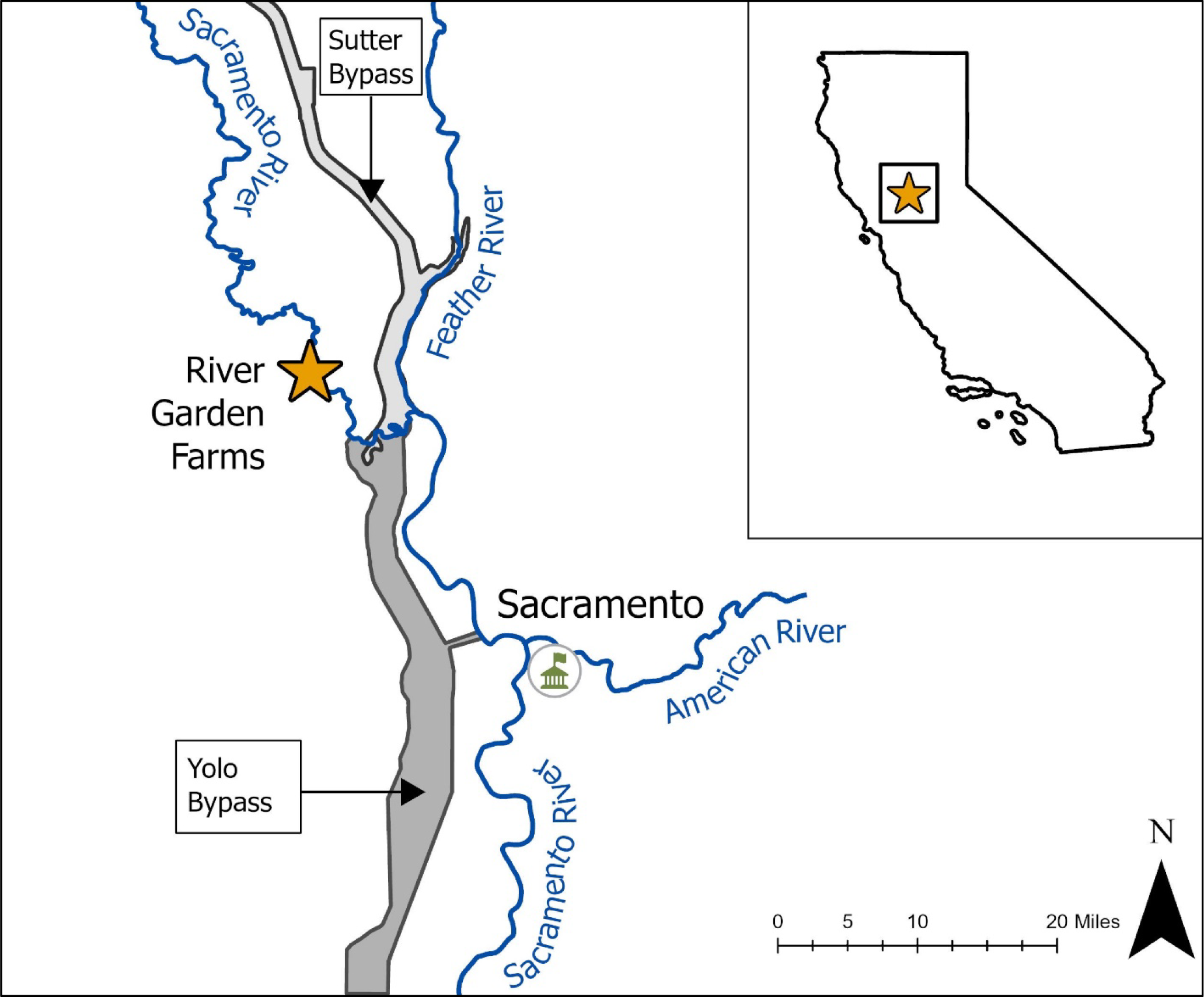
Map of study site and location of nearby flood bypasses and major rivers.

### 2.2 Experimental Plot Design

Eight 0.2 ha plots were constructed within our study site (Figure 2). We chose this location because of its proximity to a nearby surface water diversion pump which allowed for controlled flooding. We required the study field to be in rice production before and after the study to demonstrate that these methods were applicable to fields in production, and that water productivity and limnology were comparable with non-fish rearing fields. We collaborated directly with the grower and property manager through all phases of our experiment to ensure plots were structurally maintained and supplied with adequate water. Water for our plots was pumped directly from the Sacramento River into fields adjacent to the Sacramento River on the landside of the levee. These fields were called the “upper checks” where water flowed passively from one check into another through open rice boxes (structures with stacked boards to control water depth and flow rate). Water eventually flowed into our lowest check known as the “reservoir”, which supplied the water to each of the inlets of our experimental fields. The inlet and outlet of each plot was fitted with a rice box stacked with boards to maintain water depths between 0.2m-0.3m. Inlet boxes were covered in a 3-mm mesh so water could flow into the plots while preventing fish from escaping. Based on Corline et al. (2017, 2021) and Jeffres et al. (2020), we started pumping water three weeks prior to fish placement to generate longer water residence times from upstream fields into our study plots and encourage zooplankton production as a food resource for juvenile salmon.

**FIGURE 2.**
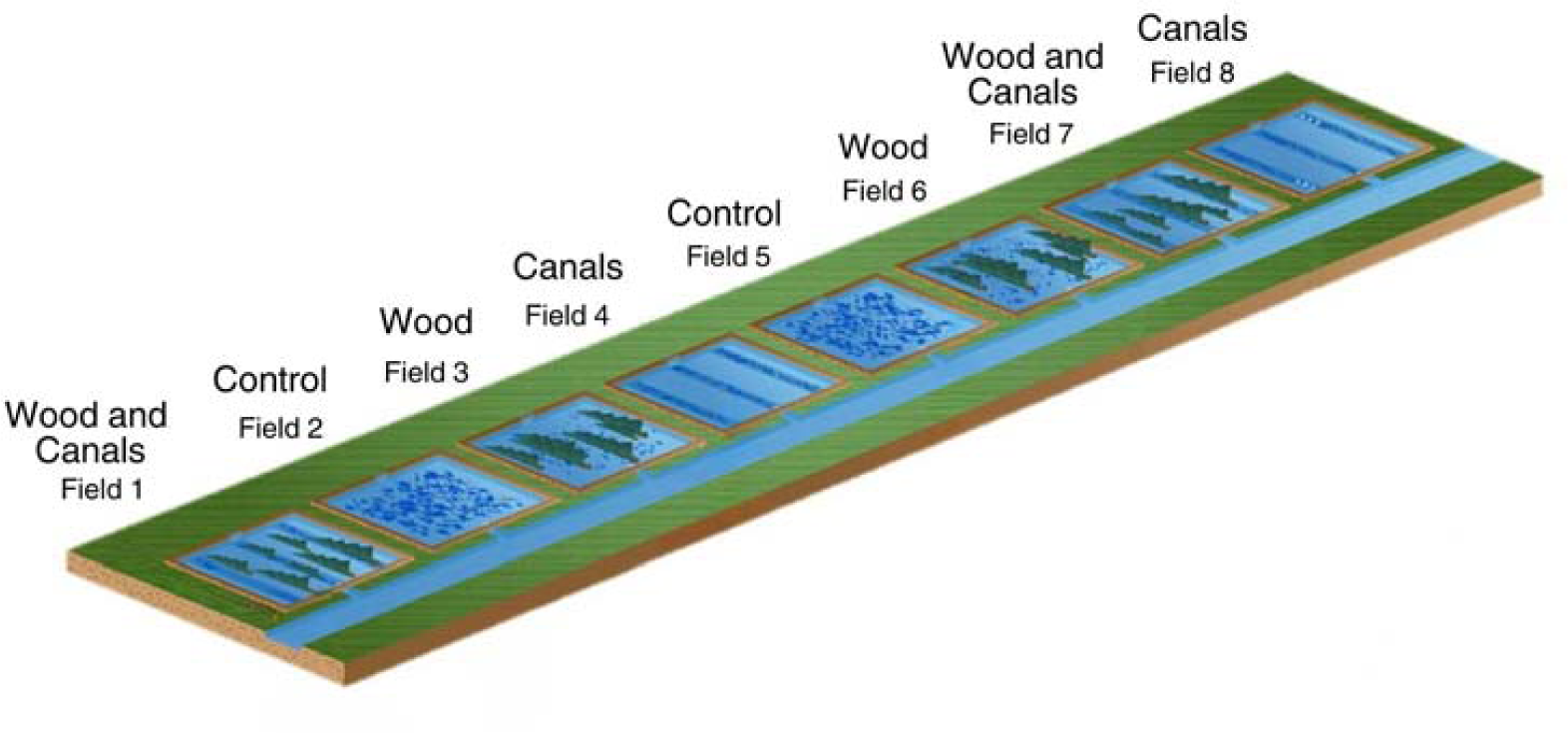
Schematic of eight experimental fields at River Garden Farms and their applied field treatments (Control, Canals, Wood, Wood and Canals). Each field contained 1000 fish with independent inflow and outflow irrigation canals.

Habitat treatment for each plot was randomly assigned by classifying each treatment as a number (1-4) and using a random number generator. Field preparation treatments were categorized into one of four treatments: large wood (wood), canals, large wood and canals (wood can canals), and control (no additional treatment). To simulate large wood on our experimental fields, we used a combination of purchased and discarded holiday trees (various spruce, fir, and pine species) because of their general availability during the study period. All plots with wood or wood and canals treatments contained 25 trees each of the following size classes and quantities: 4-5 ft (3), 5-6ft (14), 6-7 ft (5), and 7 ft+ (3). Tree sizes were not standardized due to availability. All plots had a 0.35 m deep perimeter canal to help drain the fields at the end of the study. Canal and wood and canal treatments contained 3 canals that separated the field into equal quarters from inlet to outlet. Canals were constructed to be 0.3-0.35m deep. Final canal depths varied slightly because of sediment settling after construction. Control plots received no additional treatment beyond the standard post-growing practices of chopping rice straw into stubble, baling and removing excess rice straw, and discing the field with a single pass by plowing the remaining straw into the ground to promote decomposition. All eight study fields were equipped with a HOBO U26-001 dissolved oxygen and temperature logger (Onset Computer Corporation, Bourne, Massachusetts, USA) in the center of the field that recorded conditions hourly. Loggers were secured in place using a t-post and situated 7.6 cm above the substrate. Canal and wood & canal plots had their loggers positioned 7.6 cm above the substrate of the canals to record potential temperature differences from the control plots.

### 2.3 Source and Transport of Chinook Salmon

Approximately 8,000 juvenile Chinook salmon were obtained from U.S. Fish and Wildlife Service’s Coleman National Fish Hatchery (Anderson, CA) on February 6th and 7th, 2020. To relieve osmotic stress, we created a 1.33g/L saltwater solution in our transport tank. Oxygen was supplemented into the tank through an air stone that was connected to an external oxygen tank. Dissolved oxygen (DO) and temperature were measured hourly to ensure a minimum DO concentration of 8mg/L and that water temperatures did not fluctuate more than 2°C from water sourced at the hatchery. At the study site, we added 45 mL of ammonia binding water conditioner to our transport tank to prevent nitrate toxicity as we counted and sorted fish into holding buckets. Holding buckets contained a mixture of water from the transport tank and the fields to acclimate fish to the warmer water temperatures of the study site. 1,000 fish averaging were planted within each field. Fish were not able to leave volitionally until draining commenced. Fish were not marked prior to being released on the fields due to their small body size (average fork length = 43.4mm, average mass = 0.85g). Fish were handled in strict accordance to guidelines outlined by the University of California Davis Institutional Animal Care and Use Committee (Protocol # 22685) and California Department of Fish and Wildlife Scientific Use Permit (ID # S-183530003-18360-001).

### 2.4 Field Sampling

Beginning on 17 February 2020, fish were sampled weekly using a 7.6m long beach seine net (wing mesh size 4.0 mm, center bag mesh size 3.0 mm) with a standardized effort of four pulls per plot. Each plot was seined twice prior to final draining. The first 30 fish from the first pull of each plot were sampled for length (mm ± 1) and weight (g ± 0.01). All captured fish were marked by clipping their adipose fin. Fish that measured > 50mm (∼2850 individuals over the course of the study) received a 10mm Passive Integrated Transponder (PIT) radio frequency tag (Biomark, Boise Idaho, USA).

On February 26^th^, maximum daily water temperature exceeded or approached 20°C across all the experimental fields. Due to high temperatures, we chose to conclude the study to prevent excess stress on fish. Draining began on March 2^nd^. Fish were captured in a 1.5m^3^ mesh enclosure placed into an outflow irrigation canal at the drain of each field. A 60cm Sq Low Pro Antenna IS1001 (Biomark, Boise Idaho USA) was used to count PIT tagged fish as they exited the field. Hand counts were conducted after draining to account for untagged fish. Fields were drained at night to prevent unfavorable water temperatures and reduce stress while fish exited the field. Each night, one field was drained due to the capacity of the outflow canal (8 days total).

### 2.5 Statistical Analysis

#### 2.5.1 Water Quality Across Field Treatments

To compare water temperatures across field preparation treatments, we conducted a generalized least squares (GLS) analysis using the NLME package in R (Pinheiro et al., 2023). This regression allowed us to account for correlation of temperature over time. We used a maximum likelihood estimation and specified the error term for a 2^nd^ order temporal autocorrelation error. An autocorrelation of two accounted for cyclic/diurnal dynamics found in hourly (sub-daily) measurements. Hourly temperatures were used as the response variable in our GLS model while habitat treatment was our variable of interest with our time component. We used a significance level of α= 0.05 to identify differences in temperatures across field treatments. Results from the GLS model determined if temperature would be used as fixed effect in models examining potential differences in growth and survival.

GLS Model was defined as:

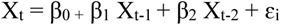

X_t_ represents a temperature estimate for t time point, β coefficient estimated by the regression, X corresponds to each of our field treatments, and ɛ_i_ is an error term to account for differences between predicted and observed temperature values.

Dissolved oxygen data were recorded on an hourly basis, however, DO measurements displayed random measurements that did not correlate with associated water temperatures. DO and temperature are known to be highly correlated with one another (Wetzel, 2001), and due to these inconsistencies, we believe our HOBO loggers were not recording DO properly and excluded these data from further analysis.

#### 2.5.2 Growth Rates

Growth data were first grouped by individual fields to account for differences in time between sampling events then by habitat treatment. Average growth rates were calculated between sampling events for each field as the difference between average fork length for each field divided by the number of days between samplings. Growth data was log transformed to meet the assumptions of normality. Assumptions of normality and homogeneity of variance were assessed using a Shapiro-Wilk and Levene test respectfully using a significance level of α < 0.05. One-way analysis of variance (ANOVA) was used to test for significant differences in growth rates across habitat treatments. Mass measurements were not included in analyses due to difficulties of collecting accurate measurements in the field.

#### 2.5.3 Survival Modeling

We used a semiparametric mixed effects Cox Proportional Hazards model (CPH; Cox, 1972) to test whether additional field treatments influenced fish survival. This approach allowed us to account for differences among individual fields by including field number as a random effect. For this model, survival was indicated as the number of fish that were counted at the end of the study as alive. To minimize strandings and drain mortalities, we employed a “fast drain” technique to maximize the outflow rate of our drain. During draining, seines and nets were used to capture any stranded fish which were then added to the cumulative survival count. Incidental mortalities at the end of draining were assumed to be caused solely by the design and implementation of draining and not habitat treatment since mortality appeared to be recent (clear eyes, silver coloring, body intact) and were always collected within the mesh enclosure in the drainage canal rather than on the fields. For these reasons all fish in the experimental fields were assumed alive at the commencement of field draining and hence survived the rearing period on the experimental fields. For our analyses, we considered these fish to have survived the duration of the experiment.

The CPH model was defined as:

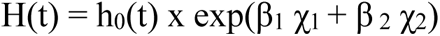

where H(t) is the estimated hazard at time t, h_0_(t) refers to the baseline hazard where it is assumed there is no effect from the covariates, β_ί_, is the regression coefficient that measures the effect size of covariates (χ_ί_). Hazard ratios (HR) were estimated for each covariate independent from time (t) based on the regression coefficients found in the CPH model HR = (exp(β_1_)). HR > 1 indicates an increase in hazard, meaning lower survival within a given habitat treatment, HR = 1 indicates no effect, HR < 1 indicates a decrease in hazard, meaning higher survival with a specific habitat treatment. CPH models were fitted using the *coxme* function in the survival analysis R package (Therneau et al., 2020). To meet the assumptions of the CPH model, proportional hazard of each covariate is assumed to be independent of time. We tested this assumption using a Schoenfeld residuals (*coxzphI*), a calculated p-value > 0.05 meets this assumption. All analyses were conducted using R v4.2.1 (R Core Team, 2022).

## 3 Results

### Temperature variations among habitat treatments

We found no statistically significant association between average hourly temperatures and habitat types (GLS: p > 0.05, Canals: p = 0.38, Wood: p = 0.15, Wood & Canals: p= 0.38, AIC= 1860.28) after accounting for AR (2) temporal correlation (Figure 2A & 2B). Daily average temperatures across treatments (9.65 - 16.58°C) were well-within accepted thermal tolerances of juvenile salmonids in the Central Valley <24°C (Figure 3A & B) (Myrick & Cech, 2001; Zillig et al., 2023). At the beginning of the study, weekly minimum temperature across all treatments averaged 4.56°C with a weekly maximum temperature of 16.37°C and a weekly average temperature between 10.13°C (Table 1). Halfway through the study (∼2 weeks later) weekly minimum temperature across treatments was 7.16°C, weekly maximum temperature was 17.61°C and a weekly average temperature of 12.55°C (Table 1). The warmest temperatures occurred during our fourth week of observations (02/24/20-03/02/20), with a weekly minimum temperature of 9.29°C, a weekly maximum temperature of 21.61°C, and a weekly average temperature of 15.59°C (Table 1). Since there was no significant difference in temperature across treatments, we did not use it as a fixed effect for our growth rate and survival analyses.

**FIGURE 3A.**
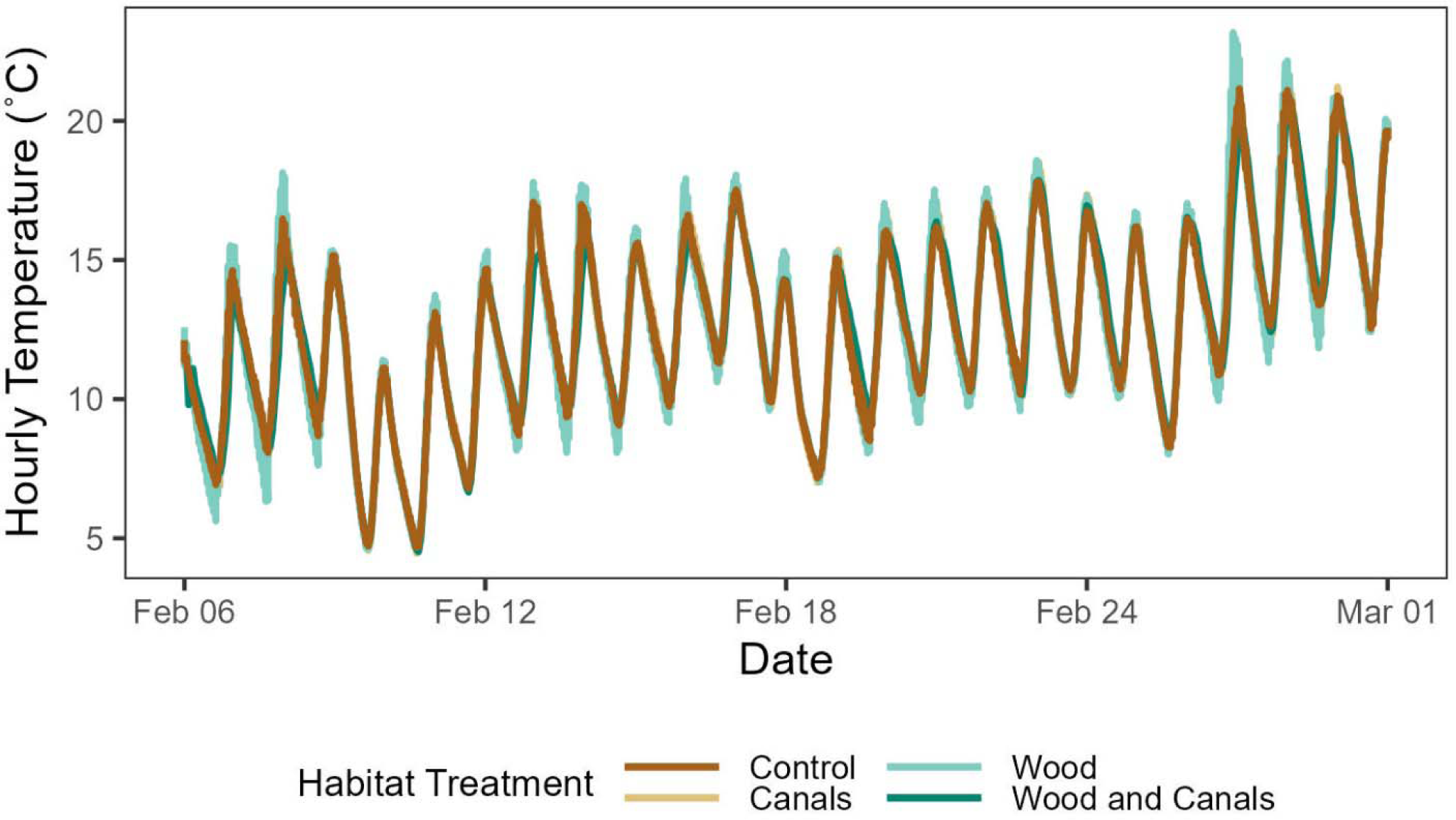
Hourly recorded temperatures from each habitat treatment: Control, Canals, Wood, Wood and Canals.

**FIGURE 3B.**
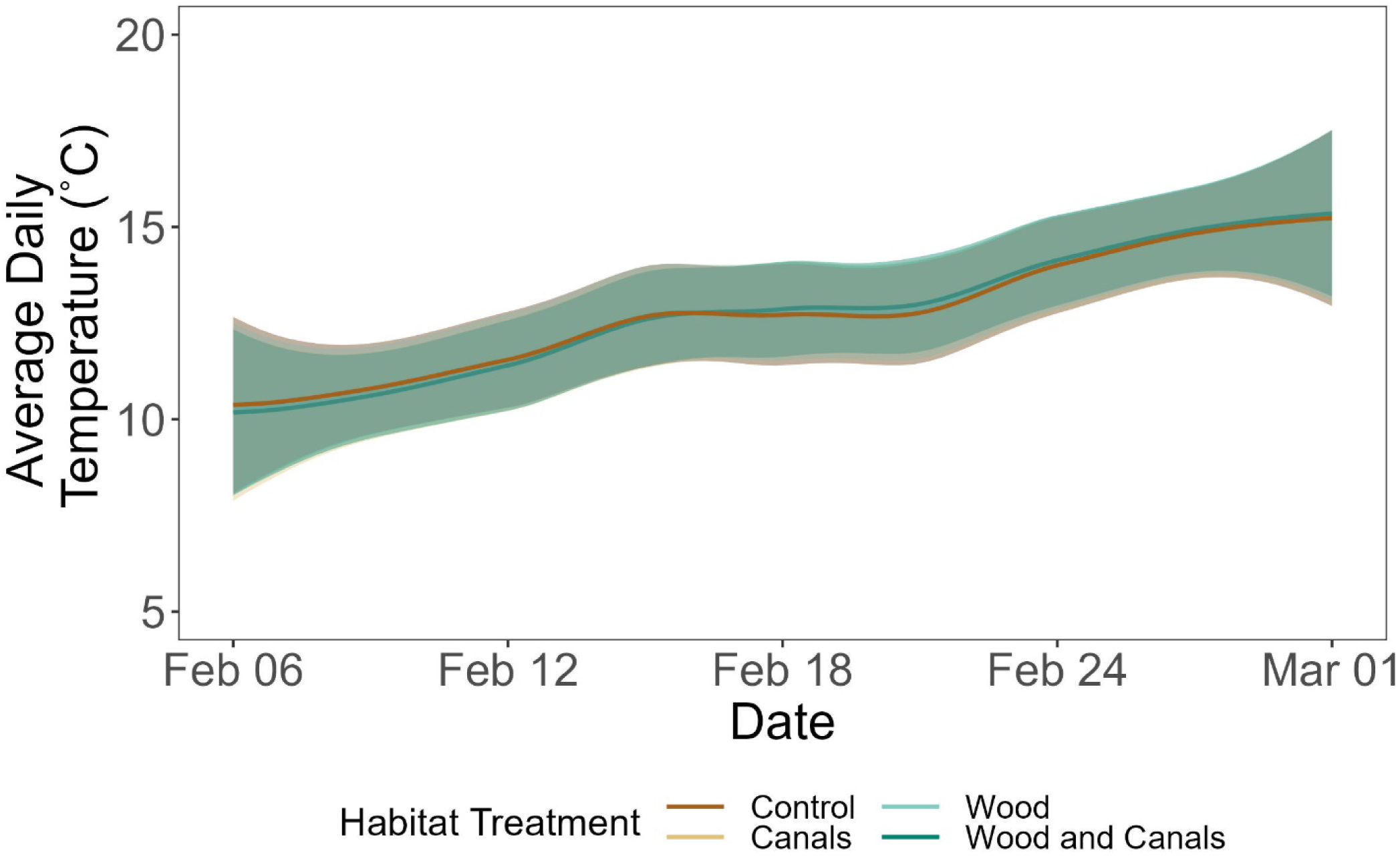
Average Daily temperatures across field treatments during the experiment using a fitted regression line. Shading represents 95% confidence interval.

**TABLE 1.** Weekly temperature summary table by habitat treatment.

| Week | Habitat Treatment | Maximum Temperature (°C) | Average Temperature (°C) | Minimum Temperature (°C) |
| --- | --- | --- | --- | --- |
| 1 | Control | 16.48 | 10.20 | 4.68 |
| 1 | Canals | 15.76 | 10.06 | 4.46 |
| 1 | Wood | 18.14 | 10.17 | 4.58 |
| 1 | Wood and Canals | 15.10 | 10.09 | 4.52 |
| 2 | Control | 17.52 | 12.59 | 7.16 |
| 2 | Canals | 17.62 | 12.53 | 7.00 |
| 2 | Wood | 18.06 | 12.60 | 7.02 |
| 2 | Wood and Canals | 17.24 | 12.49 | 7.44 |
| 3 | Control | 17.80 | 13.13 | 8.28 |
| 3 | Canals | 18.52 | 13.20 | 8.18 |
| 3 | Wood | 18.58 | 13.21 | 8.02 |
| 3 | Wood and Canals | 17.86 | 13.35 | 8.68 |
| 4 | Control | 21.16 | 15.56 | 9.00 |
| 4 | Canals | 21.28 | 15.56 | 8.98 |
| 4 | Wood | 23.18 | 15.66 | 9.70 |
| 4 | Wood and Canals | 20.80 | 15.59 | 9.46 |

### Differences in juvenile salmon growth across habitat treatments

Results from the Shapiro-Wilk test (p-value = 0.07) demonstrated that our log transformed growth rates met the assumptions of normality. However, we did not meet the assumptions of the homogeneity of variance using the Levene test (p-value = 0.04). Variance of log-transformed growth rates across treatments ranged (0.002-0.07 (log(mm)/day). Differences in variance were considered small from a biological standpoint. Results from our ANOVA showed there was no significant difference in growth rate for juvenile salmon across the different habitat treatments (ANOVA, F = 0.93, df = 3, p = 0.96). The large p-value and small variances in our growth rates support this finding. Slopes from individual linear regressions of average fork length by day for each habitat treatment resulted in estimated apparent growth rates of 0.82 mm/day for our control group, 0.83 mm/day for canals, 0.87 mm/day for wood, and 0.81 mm/day for the wood and canals group (Figure 4A & B).

**FIGURE 4.**
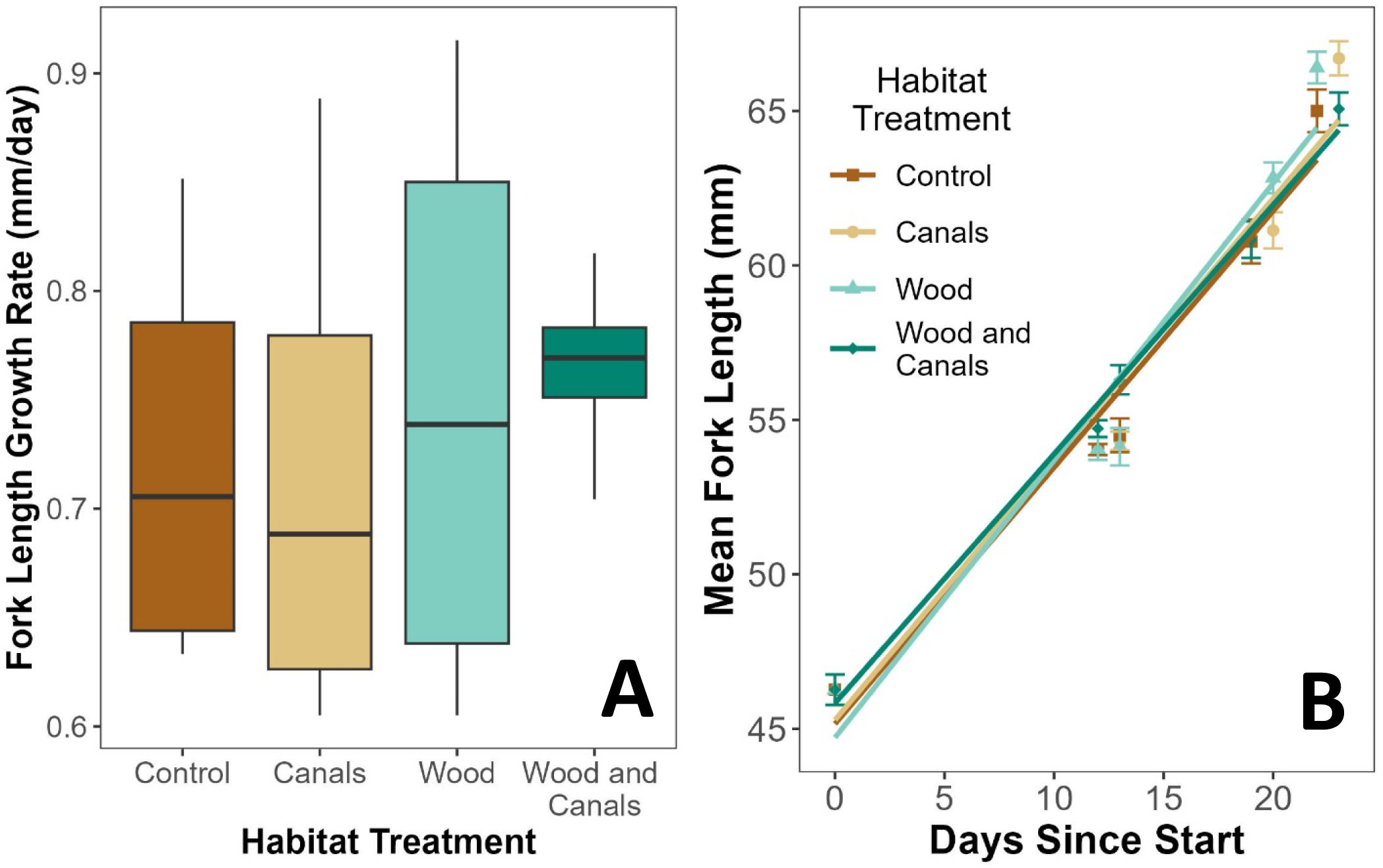
Box and whisker plot of growth rate across habitat treatments. Horizontal lines represent the median and the interquartile range, while whiskers represent the lower 25% and upper 25% of calculated growth rates. 4B: Mean fork length (mm) with standard error bars and linear regression line representing growth rate.

### Salmon Survival

In-field survival of juvenile salmon across treatments averaged 65.2% +/- 15.34%SE but ranged 50.1%-78.3% (163-839 fish) (Table 1 and Figure 5). Control treatments exhibited the highest rates of survival (78.3%). Results from our mixed effects cox proportional hazards model showed there was not a significant difference in survival across habitat treatments compared to our control group: canals (p-value = 0.30), wood (p-value = 0.97), wood & canals (p-value = 0.33) (Table 2). The variance for the random effect of field was high (229.13) with a standard deviation = 15.1. This variability may be explained by sampling events from 02/24/20-02/28/20. During that week, our study site experienced warm winter weather where air temperatures exceeded 20°C. A combination of thermal and handling stress resulted in mass mortality within one of the replicates of our canal treatment (211 morts).

**FIGURE 5.**
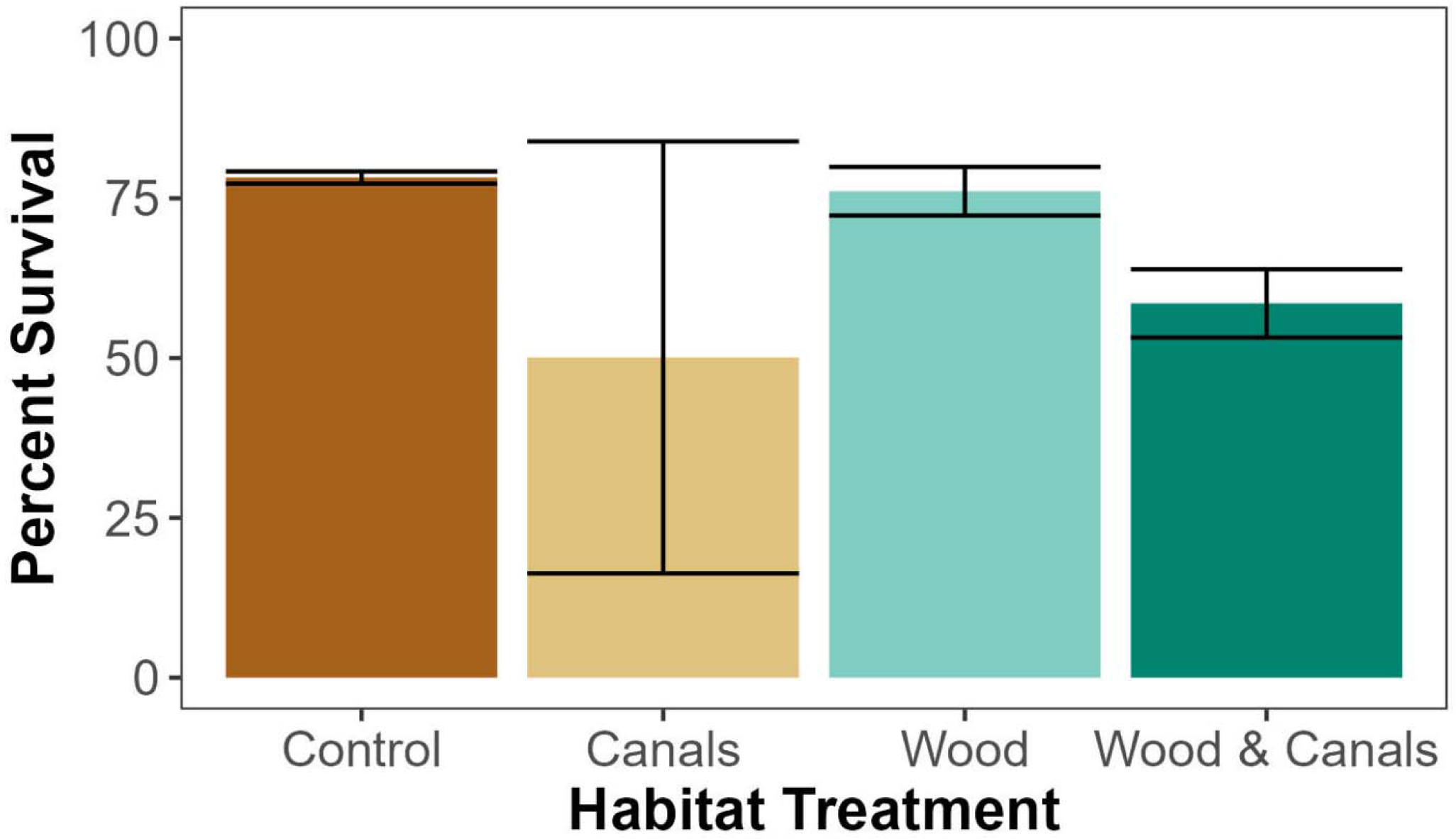
Juvenile salmon survival in habitat treatments. Bars represent mean percent survival for each treatment. Error bars denote standard error.

**TABLE 2.** Number of fish counted as alive at the conclusion of draining for each habitat treatment and their corresponding field number.

| Field | Habitat Treatment | Fish Count |
| --- | --- | --- |
| 1 | Wood and Canals | 639 |
| 2 | Control | 792 |
| 3 | Wood | 723 |
| 4 | Canals | 839 |
| 5 | Control | 773 |
| 6 | Wood | 799 |
| 7 | Wood and Canals | 532 |
| 8 | Canals | 163 |

**TABLE 3.** Cox proportional hazards model summary (AIC = 36119.22). Coefficient estimates (β) and confidence intervals (CI), Standard Error (SE) of coefficient estimates, Hazards Ratio (HR), z score (z), p-values (p) are shown below. Hazard ratios were calculated by exponentiating the β and indicates the effect of a unit of change in covariate on survival. Our control sites served as references for the categorical fixed effect of habitat treatment.

| Habitat Treatment | Coefficient ($\beta$) and CI | SE $\beta$ | HR | z | p |
| --- | --- | --- | --- | --- | --- |
| Control | -- | -- | -- | -- | -- |
| Canals | -16.17 (-46.45 to 14.11) | 15.45 | 9.51e-08 | -1.05 | 0.30 |
| Wood | -0.55 (-30.49 to 29.39) | 15.28 | 5.78e-01 | -0.04 | 0.97 |
| Wood & Canals | -15.06 (-45.22 to 15.10) | 15.39 | 2.89e-07 | -0.98 | 0.33 |

## 4 Discussion

We found increased habitat complexity through the addition of large wood and canals resulted in no significant difference in survival across treatment and control fields. The lowest levels of survival in fields at the far edges of our study site (Fields: 1,7,8). Observational photos from wildlife cameras captured piscivorous birds residing on two of these plots (Fields 7 and 8). Berms located on the outer edges of our study area (Fields 1,2,7,8) were wide enough for birds to perch and forage for fish. Fields located in the center of our study area had water eroding both sides of the berm. This resulted in narrower berms that may not have supported as much bird activity compared to perimeter berms. These observations suggest differences in survival were not the result of field treatment but rather individual field location relative to one another. Fields in the center of the study area (Fields 3,4,5,6) may have had higher fish survival because narrower berms inhibited bird perching and predation.

Compared to previous research (Holmes et al. 2021), which experienced 33.3% mortality in the first week of observations, we observed >70% survival in most of our fields (n=5) at the conclusion of our study. Holmes et al. (2021) suggest that initial mortality resulted from a combination of effects related to transport stress, acclimation to a new environment, and prey switching from pellet food to invertebrates. Transport has been shown to increase cortisol levels (an indicator of stress) in fishes, and sustained stress can result in immunosuppression and disease susceptibility (Tacchi et al., 2015). Although we do not have a measure of survival 10 days post-transfer, we may have mitigated this initial mortality by adding salt to the transport tank to minimize stress and by acclimating fish to their new environment by adding field water into holding buckets prior to release.

All fields contained a perimeter trench that surrounded the edges of the field regardless of field treatment. Similar to Holmes et al. (2021), we suggest perimeter trenches acted as migratory corridors that encouraged fish to emigrate off the fields during draining and resulted in reducing the occurrence of strandings. Our study supports the findings of Ogaz et al. (2022) and Holmes et al. (2021), which found that fish strandings are reduced, and passage facilitated, where there is high out-flow. The fast drain technique may have triggered a sensory que for fish to emigrate off the fields, resulting in a lower number of observed strandings. Based on our results and the findings of Holmes et al. (2021) we suggest the inclusion of perimeter trenches in agricultural floodplain fish rearing practices.

Towards the end of our study, we observed water temperatures that exceeded 20°C. Previous research on juvenile Chinook Salmon in the Central Valley found optimal growth occurs between 17-20°C (Zillig et al., 2021), however issues arise when juvenile Chinook are exposed to temperatures exceeding 20°C for extended periods of time. Marine & Cech Jr (2004) found that juvenile Chinook chronically exposed to temperatures between 21-26°C in the laboratory exhibited slowed growth rates, decreased appetites, impaired smoltification, and were more vulnerable to predation. Under these conditions, 21-26°C reared fish were smaller compared to cohorts of fishes reared at 13-16°C and 17-20°C. Marine & Cech Jr (2004) suggest rearing juvenile Chinook in temperatures >20°C may result in sublethal effects that impair swim performance. While our fish were exposed to temperatures >20°C, it is important to note these data depict the diurnal temperature fluctuations of the day. Temperatures that exceeded 20°C were typically recorded during the warmest time of the day and decreased soon after. Although we experienced massive mortality in one of our canal replicates during a week of warm weather, this most likely resulted from a combination of thermal and handling stress. While water temperatures quickly returned to a preferable range, potential adopters of this practice should be aware of the adverse water conditions that arise when air temperatures exceed 20°C.

Calculated growth rates from our experimental fish suggest additional field treatments were not needed to support high growth rates in juvenile Chinook Salmon. Our calculated mean growth rates of 0.81 - 0.87mm/day were similar to those found in previous studies on California agricultural floodplains (0.70 mm/day +/- 0.01 mm/day) (Holmes et al. 2021) and high compared to previous growth rates of river-reared fish (0.43 mm/day +/- 0.03 mm/day – 0.52mm/day +/- 0.02mm/day) (Sommer et al., 2001). Habitat conditions that promote high growth rates result in larger body sizes which have been found to increase fitness in most fish species (Ebersole et al., 2006; Hayes et al., 2008; McCormick et al., 1998; Sundström et al., 2005; Unwin, 1997). Growing salmon on agricultural floodplains may support native salmon populations by increasing their out-migration survival through high growth rates during their rearing stage.

Although this study helped answer some critical questions regarding juvenile salmon on agricultural floodplains, there are several notable limitations. One factor that could have affected our results is a thiamine deficiency found in most Central Valley salmon beginning in broodstock year 2019. Thiamine Deficiency Complex (TDC) was first observed in Central Valley Chinook in the winter of 2020 from recently hatched juveniles that had low levels of thiamine (Suffridge et al. 2023). Yancheff et al. (2023, Preprint) found that up to 50% of eggs sampled from Coleman National Fish Hatchery fell below the minimum 5 nmol/g needed for fry viability. Thiamine is an essential micronutrient fish use to metabolize fatty acids into energy (Bell 2022, Preprint). Fish deficient in thiamine exhibit chaotic swim patterns (e.g., spiraling), loss of equilibrium, lethargy, and impaired neurological functions (Bell 2022, Preprint). If left untreated, it can lead to high mortality in juvenile fish (Bell 2022, Preprint; Yancheff 2023, Preprint; Suffridge et al. 2023). While Coleman National Fish Hatchery immediately began thiamine baths to mitigate this deficiency, there is no confirmation that our study fish were treated for thiamine prior to release.

A limitation to our study design was working on constructed plots rather than using established fields. To meet the objectives of our study, we needed small fields that we could easily change the habitat conditions. This resulted in a low number of replicates of constructed fields. Low replication requires a strong difference in both replicates to signal a significant difference in either growth or survival across our treatments. However, our calculated p-values for growth rate and (ANOVA p-value= 0.962) and survival (Cox Proportional Hazards: canal: p-value = 0.3, wood: p-value = 0.97, wood & canals: p-value = 0.33) suggest there were no significant differences in either response variable across treatments despite low replication. However, our low number of replicates may explain why we observed a high standard error in our cox proportional hazards model. Our canal treatment had the largest variance in survival because one replicate had the highest number of fish (n=839) survive while the replicate field had the lowest number (n=163).

Another constraint arose from our interest in demonstrating the applicability of fish rearing during the non-growing season. We chose to complete this study outside of nearby flood bypasses, known as the “dry side”, to ensure the study was not comprised by flooding. There were a limited number of fields that met these requirements and were easily accessible. Our study location required us to manually flood our fields which created homogenous flow conditions. In contrast, fields within the flood bypasses experience ephemeral flood conditions with dramatic fluctuations in flow during the winter and early spring (0 - > 4,000m^3^/sec) (Sommer et al. 2004). Small flood events cause localized flooding within the bypass where water is sourced from nearby creeks and sloughs and is hydrologically separated into distinct segments along the bypass (Sommer et al., 2001). During large flood events, the flood bypasses’ main source of water come from flows redirected from the Sacramento and Feather Rivers. Different sources of water influence the macroinvertebrate assemblages found within these fields and potentially influences fish growth. Sommer et al. (2004) found during high flow events, bypass fields exhibited high concentrations of *Chironomids* while Corline et al. (2017) found a higher concentration of *Daphniids* during non-flood years and recorded some of the highest growth rates of juvenile Chinook Salmon in the Central Valley. Manually flooding the fields meant we used one source of water with consistent slow-moving conditions. While this may favor juvenile salmon foraging, these homogenous conditions do not mimic the ephemeral nature of a natural floodplain. Conducting the study on the dry side with manual flooding allows for flood control at the expense of excluding predators from entering the study site. Future studies should consider the limitations/benefits to manually flooding the fields and how it affects the predator prey dynamics between juvenile salmon and aquatic predators.

Overall, our study supports previous research that winter-flooded agricultural floodplains support high growth and survival of juvenile salmon. During the winter, juvenile salmon typically move downstream and use off-channel floodplains to rear and forage as part of their out-migration (Kjelson et al., 1982). In natural floodplains, salmon migrate off floodplains into the mainstem river to start their ocean migration. In contrast, rice production occurs during the mostly dry growing season, mid-spring through fall. When combined, these two timelines would allow resource managers to apply a novel management approach by utilizing agricultural fields for the purpose of promoting a native fish species. Parallel conservation programs have been highly successful for migratory birds and waterfowl (Bird et al., 2000; Elphick & Oring, 1998; Strum et al., 2013; USA Rice, 2018). A potential agricultural practice standard for fish (e.g., through the USDA NRCS program) would encourage farmers to flood their fields for the benefit of native fishes while maintaining them for production during the growing season.

Across the Pacific Rim, particularly in California, salmon populations continue to decline amongst a highly modified and human dominated landscape (Brown et al., 2011; Moyle, 2002; Nehlsen et al., 1991; Yoshiyama et al., 1998). The loss of wetland habitat has been identified as a major contributor to a decline in Central Valley salmonids (Mount, 1995; Tockner & Stanford, 2002; Williams et al., 2009). This is expected, considering that these species evolved on a landscape dominated by seasonally inundated wetlands that varied in response to a Mediterranean climate (Cordoleani et al., 2021). To prevent these species from continued decline, novel conservation approaches are needed to increase salmon resiliency (Dudgeon et al., 2006; Moyle, 2014; Rosenzweig, 2003a, 2003b). Within California, access to floodplain habitats is critical to improving growth and fitness of juvenile Chinook Salmon populations (Sommer et al. 2001b; Jeffres et al. 2008; Katz et al. 2017; Goertler et al. 2018; Holmes et al. 2021). However, management actions should prioritize making floodplain habitat more accessible and beneficial for juvenile salmon. Future work may focus on increasing connectivity between the main river channel and nearby flood bypasses (Bayley, 1991; Jeffres et al., 2008; Junk et al., 1989; Moyle et al., 2007), incorporating functional flows by managing upstream dams’ water retention and release (Kiernan et al., 2012; Yarnell et al., 2010, 2015, 2020), assessing what current management practices favor poor water quality or predation (Bennett, 2005; Crain et al., 2004; Moyle, 2002; Moyle et al., 2011; Yarnell et al., 2010), and examining how increasing floodplain habitat may benefit other native fish species (Crain et al., 2004; Feyrer et al., 2006; Moyle et al., 2007).

While agricultural floodplains are not the same as natural wetlands, this study and others (Corline et al., 2017; Holmes et al., 2021; Jeffres et al., 2020; Katz et al., 2017; Sommer et al., 2001a; Sommer et al., 2001b) generally supports the idea that agricultural floodplains offer critical benefits to native fishes. It is our hope that these data be used to inform agricultural practices and conservation policies that would encourage farmers to manage their fields to promote the recovery of California salmon. Furthermore, application of reconciliation frameworks may have similar potential in other parts of the world where large river floodplains and agricultural production intersect. Finding methods that simultaneously support food production and freshwater biodiversity will be essential to protecting freshwater biodiversity crisis while supporting human populations.

## ACKNOWLEDGEMENTS

We especially thank Paul Buttner with the California Rice Commission for logistical support and stakeholder engagement. We thank Amanda Agosta, Heather Bell, Mackenzie Miner, Hailey Gleason, Adrian Loera, Wilson Xiong and additional volunteers from the Fish Conservation Physiology and Rypel Labs for their tremendous help in conducting fieldwork and collecting data. A major thank you to River Garden Farms, especially Roger Cornwell and Dominic Bruno for assisting in finding suitable fields for this experiment and collaborating with our team on project design and adaptive water management of the fields – this project would not have been possible without their full support. We also thank Jacob Katz, Jacob Montgomery, and Jennifer Kronk at California Trout for their feedback and guidance. We would also like to thank California Department of Fish & Wildlife for their input on project design and implementation as well as coordination of the scientific collecting permit.

## DATA AVAILABILITY STATEMENT

All datasets associated with this study are published as part of this paper.

## ETHICS APPROVAL STATEMENT

All animal housing and experiments were conducted in strict accordance with the UC Davis Institutional Animal Care & Use Committee (IACUC) Protocol # 22685.

